# Using Local Genetic Correlation Improves Polygenic Score Prediction Across Traits

**DOI:** 10.1101/2022.03.10.483736

**Authors:** Oliver Pain, Cathryn M. Lewis

**Author notes:** <u>Corresponding author:</u> Oliver Pain.

## Abstract

**Introduction:** The predictive utility of polygenic scores (PGS) is steadily increasing as genome-wide association studies (GWAS) increase in sample size and diversity, and as PGS methodology is further developed. Multivariate PGS approaches incorporate GWAS results for secondary phenotypes which are genetically correlated with the target phenotype. These improve prediction over using PGS for only the target phenotype. However, previous methods have only considered the genome-wide estimates of SNP-based heritability (*h*^2^_SNP_) and genetic correlation (*r*_*g*_) between target and secondary phenotypes. In this study, we assess the impact of local *h*^2^_SNP_ and *r*_*g*_ within specific loci on cross-trait prediction.

**Methods:** We evaluate PGS using three target phenotypes (depression, intelligence, BMI) in the UK Biobank, with GWAS summary statistics matching the target phenotypes and 14 genetically correlated secondary phenotypes. PGS SNP-weights were derived using MegaPRS. Local *h*^2^_SNP_ and *r*_*g*_ were estimated using LAVA. We then evaluated PGS after reweighting SNP-weights according to local *h*^2^_SNP_ and *r*_*g*_ estimates between the target and secondary phenotypes. Elastic net models containing PGS for multiple phenotypes were evaluated using nested 10-fold cross validation.

**Results:** Modelling target and secondary PGS significantly improved target phenotype prediction over the target PGS alone, with relative improvements ranging from 0.8-12.2%. Furthermore, we show reweighting PGS by local *h*^2^_SNP_ and *r*_*g*_ estimates can enhance the predictive utility of PGS across phenotypes, with additional relative improvements of 0.2%-2.8%. Reweighting PGS by local *h*^2^_SNP_ and *r*_*g*_ improved target phenotype prediction most when there was a mixture of positive and negative local *r*_*g*_ estimates between target and secondary phenotypes.

**Conclusion:** Modelling PGS for secondary phenotypes consistently improves prediction of target phenotypes, and this approach can be further enhanced by incorporating local *h*^2^_SNP_ and *r*_*g*_ estimates to highlight relevant genetic effects across phenotypes.

## Introduction

The predictive utility of polygenic scores (PGS) is steadily increasing as genome-wide association studies (GWAS) increase in sample size and become available for a wider range of phenotypes and populations. Furthermore, advances in PGS methodology are making important improvements to the predictive utility of PGS (1). PGS methods that utilise GWAS summary statistics together with linkage disequilibrium (LD) reference data typically provide the highest predictive utility as individual-level data for large sample sizes is often unavailable.

PGS are a commonly used research tool and are being increasing studied for clinical application to enhance personalised medicine. PGS for a target phenotype typically only explain a small proportion of variance and will therefore be most useful when integrated into prediction models that also consider other predictors. For example, integrating coronary artery disease PGS with established clinical predictors significantly improved prediction of coronary artery disease events (2). Prediction of a target phenotype can also be improved by incorporating PGS for secondary phenotypes that are genetically correlated with the target phenotype (3).

Other multivariate approaches have also been considered for improving the prediction of a target phenotype, such as SMTpred (4), Genomic SEM (5), and MTAG (6). All these approaches improve prediction of a target phenotype by incorporating GWAS summary statistics for genetically correlated secondary phenotypes. However, these previous approaches only consider the genome-wide genetic correlation (*r*_*g*_) between the target and secondary phenotypes, thereby not allowing for *r*_*g*_ to differ at specific loci. Recently developed methods allow the estimation of local *r*_*g*_, such as LAVA (7) and HESS (8). This more granular insight into the shared and unique genetic effects across traits has highlighted that although two phenotypes may have a significant genome-wide *r*_*g*_, this may be driven by only a few loci, and may even involve a mixture of positive and negative *r*_*g*_ loci. Current multivariate PGS approaches assume a consistent *r*_*g*_ across the genome and will therefore incorporate genetic effects from secondary phenotypes that are irrelevant and potentially inversely related to the target phenotype, thereby reducing the value of adding the secondary PGS in prediction.

In this study we explore whether local *r*_*g*_ estimates can be leveraged to enhance the predictive utility of PGS across phenotypes. In a similar approach to Krapohl et al., we use an elastic net to model PGS for the target PGS and multiple secondary PGS (3). We then evaluate the effect of reweighting variants within specific loci based on local *r*_*g*_ estimates computed using LAVA. Using three traits of depression, intelligence and BMI in the UK Biobank, our findings support previous literature showing improved prediction of the target phenotype prediction when modelling PGS for secondary phenotypes over target phenotype PGS alone. Further, we demonstrate for the first time that using local *r*_*g*_ alongside the secondary PGS provides further statistically significantly improvements in predicting the target phenotype.

## Methods

### UK Biobank (UKB)

UKB is a prospective cohort study that recruited >500,000 individuals aged between 40-69 years across the United Kingdom (9). The UKB received ethical approval from the North West -Haydock Research Ethics Committee (reference 16/NW/0274).

#### Phenotype data

Three UKB phenotypes were analysed: depression, body mass index (BMI) and intelligence.

For depression, UKB participants were coded as cases if they met the Composite International Diagnostic Interview Short Form criteria for lifetime depression which was assessed in the online Mental Health Questionnaire (MHQ) using scoring protocols proposed by Davis et al (10). Depression cases were screened for indications of schizophrenia or bipolar disorder in the MHQ. Controls were excluded if they showed any psychiatric indications in the MHQ or other depression indications: ICD-10 diagnoses; endorsement of self-reported depression; endorsement of current antidepressant usage; single or current depression according to the Smith criteria (11). Full details of the exclusion criteria have been previously described (12).

BMI was defined using the Body mass index variable (Field ID: f.21001.0.0).

Intelligence was defined using the Fluid intelligence score, assessed using the 13 item UKB Touch-screen Fluid intelligence test (13). The test measures the capacity to solve problems that require logic and reasoning ability, independent of acquired knowledge. The fluid intelligence variable (Field ID: f.20191.0.0) was derived by UKB as an unweighted sum of the number of correct answers, assigning a score of 0 to unanswered questions.

This study analysed a subset of ∼50,000 UKB participants for each phenotype to reduce computation burden. For intelligence and BMI, a random sub-sample was selected. For depression, a random sample of 25,000 cases and 25,000 controls was selected.

#### Genetic data

UKB released imputed dosage data for 488,377 individuals and ∼96 million variants, generated using IMPUTE4 software (9) with the Haplotype Reference Consortium reference panel (14) and the UK10K Consortium reference panel (15). This study retained individuals that were of European ancestry based on 4-means clustering on the first two principal components provided by the UKB, had congruent genetic and self-reported sex, passed quality assurance tests by UKB, and removed related individuals (>3^rd^ degree relative, KING threshold > 0.044) using relatedness kinship (KING) estimates provided by the UKB (9). The imputed dosages were converted to hard-call format for all variants.

### Polygenic scoring

Polygenic scores were derived within a reference-standardised framework, where polygenic scores are derived using a common set of genetic variants, linkage disequilibrium estimates, and allele frequency estimates (1). This reproducible and standardised approach is good research practice and is also well suited for the clinical setting.

#### SNP-level QC

HapMap3 variants from the LD-score regression website (see Web Resources) were extracted from UKB, inserting any HapMap3 variants that were not available in the target sample as missing genotypes (as required for reference MAF imputation by the PLINK allelic scoring function) (16). No other SNP-level QC was performed.

#### Individual-level QC

Individuals of European ancestry were retained for polygenic score analysis. They were identified using 1000 Genomes Phase 3 projected principal components of population structure, retaining only those within three standard deviations from the mean for the top 100 principal components. This process will also remove individuals who are outliers due to technical genotyping or imputation errors.

#### GWAS summary statistics

GWAS summary statistics independent of UK Biobank were identified for the three target phenotypes, and 14 secondary phenotypes previously shown to have a genetic correlation with at least one of the target phenotypes (Table 1). GWAS summary statistics underwent quality control to extract HapMap3 variants, remove ambiguous variants, remove variants with missing data, flip variants to match the reference, retain variants with a minor allele frequency (MAF) > 0.01 in the European subset of 1KG Phase 3, retain variants with a MAF > 0.01 in the GWAS sample (if available), retain variants with a INFO > 0.6 (if available), remove variants with a discordant MAF (>0.2) between the reference and GWAS sample (if available), remove variants with association p-values >1 or </=0, remove duplicate variants, and remove variants with sample size >3SD from the median sample size (if per variant sample size is available).

**Table 1.** GWAS summary statistics, for three target phenotypes, depression, body mass index and intelligence and 14 secondary phenotypes.

| <u>Code</u> | <u>Phenotype</u> | <u>N</u> | <u>Ncase</u> | <u>Ncon</u> | <u>Prev.</u> | <u>PMID</u> | <u>Ref.</u> |
| --- | --- | --- | --- | --- | --- | --- | --- |
| *Target phenotypes* |  |  |  |  |  |  |  |
| DEPR06 | Major Depression (excl. UKB) | 431394 | 116404 | 314990 | 0.15 | 29700475 | (24) |
| INTE03 | Intelligence | 269867 | NA | NA | NA | 29942086 | (25) |
| BODY04 | BMI | 322154 | NA | NA | NA | 25673413 | (26) |
| *Secondary phenotypes (binary)* |  |  |  |  |  |  |  |
| SCHI02 | Schizophrenia | 77096 | 33640 | 43456 | 0.01 | 25056061 | (27) |
| BIPO02 | Bipolar Disorder | 51710 | 20352 | 31358 | 0.015 | 31043756 | (28) |
| AUTI07 | Autism | 46350 | 18381 | 27969 | 0.012 | 30804558 | (29) |
| SMOK04 | Smoking (Ever/Never Regular) | 74035 | 41969 | 32066 | 0.5 | 20418890 | (30) |
| ADHD05 | Attention Deficit Hyperactivity Disorder | 53293 | 19099 | 34194 | 0.05 | 30478444 | (31) |
| DIAB05 | Type-2 Diabetes | 159208 | 26676 | 132532 | 0.05 | 28566273 | (32) |
| OBES01 | Obesity Class 1 | 98697 | 32858 | 65839 | 0.13 | 23563607 | (33) |
| COAD01 | Coronary Artery Disease | 184305 | 60801 | 123504 | 0.03 | 26343387 | (34) |
| *Secondary phenotypes (continuous)* |  |  |  |  |  |  |  |
| ANXI02 | Anxiety (Factor Score) | 17310 | NA | NA | NA | 26754954 | (35) |
| MENA01F | Age at Menarche | 132989 | NA | NA | NA | 25231870 | (36) |
| WAIS01 | Waist Circumference | 231927 | NA | NA | NA | 25673412 | (37) |
| GLYC05 | Fasting Insulin (Age- & Sex-adjusted) | 38238 | NA | NA | NA | 20081858 | (38) |
| INTE01 | Childhood Intelligence | 12441 | NA | NA | NA | 23358156 | (39) |
| COLL01 | College Completion | 95427 | NA | NA | NA | 23722424 | (40) |

*Note. Prev. = Assumed population prevalence for binary GWAS phenotypes; Ref. = Reference for each GWAS.*

#### Reference genotype datasets

Target sample genotype-based scoring was standardised using the European subset of 1000 Genomes Phase 3 (N=503).

#### MegaPRS

GWAS summary statistics were processed for polygenic scoring using MegaPRS (17), as implemented by LDAK. MegaPRS implements polygenic scoring approaches using the LDAK heritability model (18), is computationally efficient, and has good predictive utility compared with other widely-used PRS methods (1)(See URLs for updated results including MegaPRS). Like many PGS methods, MegaPRS uses a range of effect size distribution parameters to optimise the PGS. For simplicity, we used the model selected using the pseudo summary approach, also referred to as pseudo validation approach, which estimates the best set of parameters without requiring an external validation sample. This has been shown to perform well compared to other pseudo validation approaches and performs similarly to the best models identified using formal validation procedures.

#### LAVA

LAVA (Local Analysis of [co]Variant Annotation) was used to estimate the local *h*^2^_SNP_ and *r*_*g*_ between target and secondary phenotypes (7). LAVA splits the genome into 2,495 non-overlapping and broadly LD independent loci, and then for each locus LAVA estimates the SNP-based heritability for a given phenotype, and the genetic correlation between phenotypes. As recommended, the genetic covariance intercept estimated by bivariate LD score regression was used to account for any sample overlap between GWAS (19). LAVA was then run between each target GWAS and all secondary GWAS using the run.univ.bivar function, with default settings, restricting the bivariate *r*_*g*_ test to loci with a *h*^2^_SNP_ p-value < 0.05 for both phenotypes. When running LAVA, all GWAS were set as continuous phenotypes to reduce computation time, and local *h*^2^_SNP_ estimates were subsequently converted to the liability scale for binary phenotypes (assumed population prevalence listed in Table 1) (20).

#### Reweighting PGS by local h^2^_SNP_ and r_g_

Several approaches were used to reweight SNPs -with local *h*^2^_SNP_ and *r*_g_ estimates from LAVA. First, we restricted all PGS analyses to a subset of SNPs with local *h*^2^_SNP_ p-value < 0.05 for both phenotypes. We then further defined subsets of these SNPs with local *r*_*g*_ p-value < 0.05, and with false discovery rate (FDR)-corrected local *r*_*g*_ p-value < 0.05. These thresholds give three SNP subsets of increasing stringency, where the numbers of loci included within the reweighted PGS, depend on the statistical significance of local *h*^2^_SNP_ and *r*_*g*_ estimates. We then reweighted variants in MegaPRS using two approaches scaling the secondary phenotype PGS effect size (β) by (1) local *r*_*g*_ estimates only, *β*_*secondary*_ × local *r*_*g*_, and 2) local *r*_*g*_ estimates scaled by the ratio of local *h*^2^_SNP_ between target and secondary phenotypes, *β*_*secondary*_ × *local r*_*g*_ × *h*^*2*^_*SNP-target*_ / *h*^*2*^_*SNP-secondary*_.

After GWAS summary statistics were processed by MegaPRS and reweighted according to LAVA estimates, polygenic scores were calculated using PLINK with reference MAF imputation of missing data (16). All scores were standardized (scaled and centred) based on the mean and standard deviation of polygenic scores in the reference sample.

#### Evaluating PGS

Prediction accuracy was evaluated as the Pearson correlation between the observed and predicted phenotype outcomes. Correlation was used as the main test statistic as it is applicable for both binary and continuous phenotypes, and standard errors are easily computed. Correlations can be converted to test statistics such as *R*^*2*^ (observed or liability) and area under the curve (AUC) (equations 8 and 11 in (20)), with relative performance of each method remaining unchanged.

Logistic regression was used for predicting binary phenotypes, and linear regression for predicting continuous phenotypes. If the model contained only one predictor, a generalized linear model was used. If the model contained more than one predictor, an elastic net model was applied to avoid overfitting from including multiple correlated predictors (21).

A nested cross validation procedure (22) was used to estimate the predictive utility of each model, where hyperparameter selection is performed using inner 10-fold cross-validation, while an outer 5-fold cross-validation computes an unbiased estimate of the predictive utility of the model with the inner cross-validation based hyperparameter tuning. This approach avoids overfitting whilst providing modelling predictions for the full sample. The inner 10-fold cross validation for hyperparameter optimisation was carried out using the ‘caret’ R package.

The correlations between observed and predicted values of each model were compared using the William’s test (also known as the Hotelling-Williams test) (23) as implemented by the ‘psych’ R package’s ‘paired.r’ function, with the correlation between model predictions of each method specified to account for their non-independence. A two-sided test was used when calculating *p*-values.

## Results

PGS for three target phenotypes and 14 secondary phenotypes were derived using MegaPRS, and SNPs were reweighted with local *h*^2^_SNP_ and *r*_*g*_ estimates. Models using nested 10-fold cross-validation were then derived to evaluate whether secondary PGS can improve target phenotype prediction over the target PGS alone, and whether reweighting secondary PGS by local *h*^2^_SNP_ and *r*_*g*_ estimates can further improve target phenotype prediction. See Figure 1 for a schematic representation this study design.

**Figure 1.**
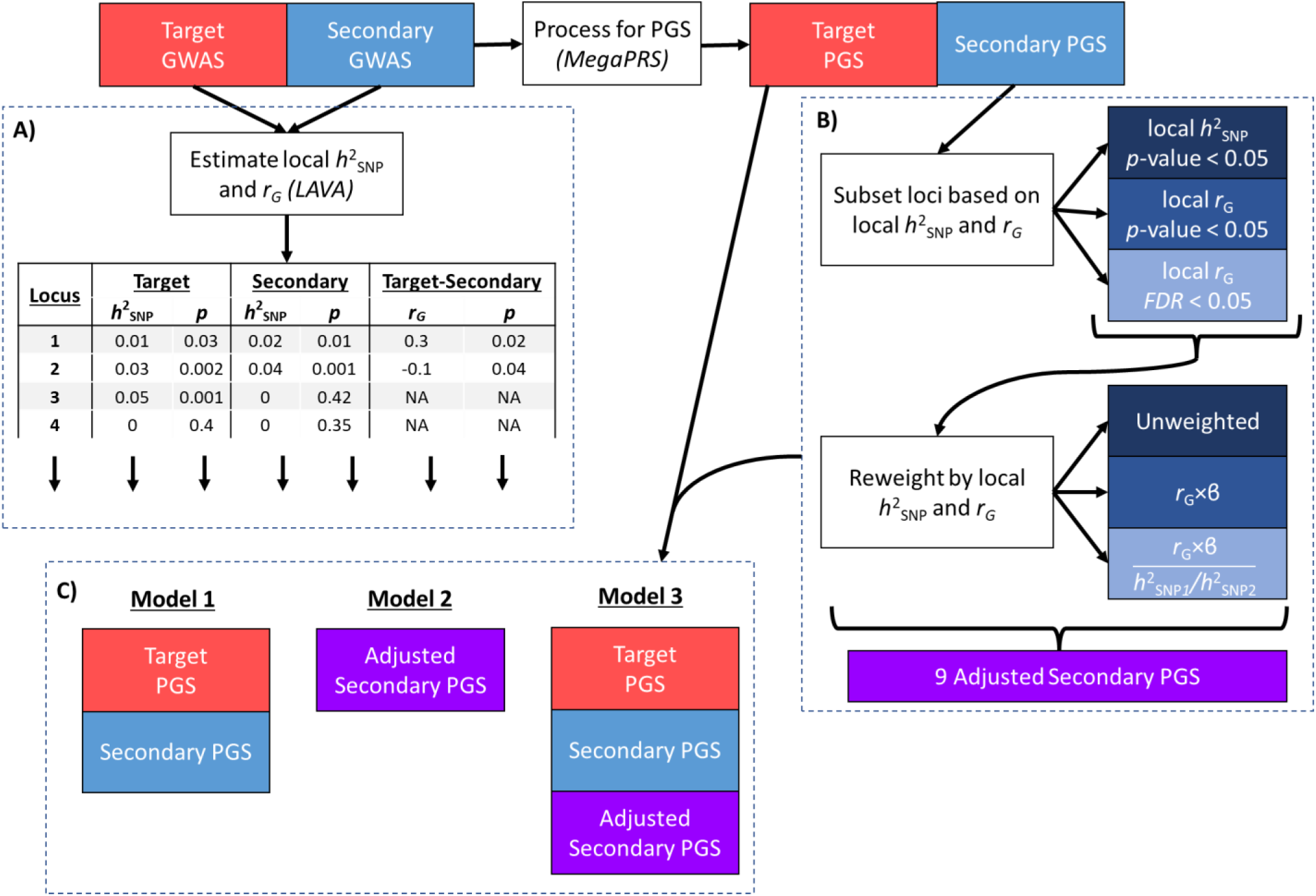
Schematic representation of study design. A) Local h^2^_SNP_ and r_g_ are estimated for target and secondary GWAS. B) After processing by MegaPRS, PGS are stratified and reweighted according to local h^2^_SNP_ and r_g_. C) Models containing standard PGS, and local h^2^_SNP_ and r_g_ adjusted PGS are evaluated and compared.

Genome-wide estimates of *h*^2^_SNP_ and *r*_*g*_ from LD score regression supported our selection of target and secondary GWAS, with statistically significant estimates of *h*^2^_SNP_ for all phenotypes, and at least 13 statistically significant estimates of *r*_*g*_ between target and secondary phenotypes (Figure 2). In local analysis, LAVA often highlighted a mixture of positive and negative local *r*_*g*_ estimates between target and secondary phenotypes, even where the genome-wide genetic correlation was not significant (Figure 2). For example, the genome-wide *r*_*g*_ between major depression (DEPR06) and adult intelligence (INTE03) was not statistically significant (*r*_*g*_ = −0.013, SE = 0.024), but LAVA identified 43 positive and 57 negative local genetic correlations surviving FDR correction for multiple testing.

**Figure 2.**
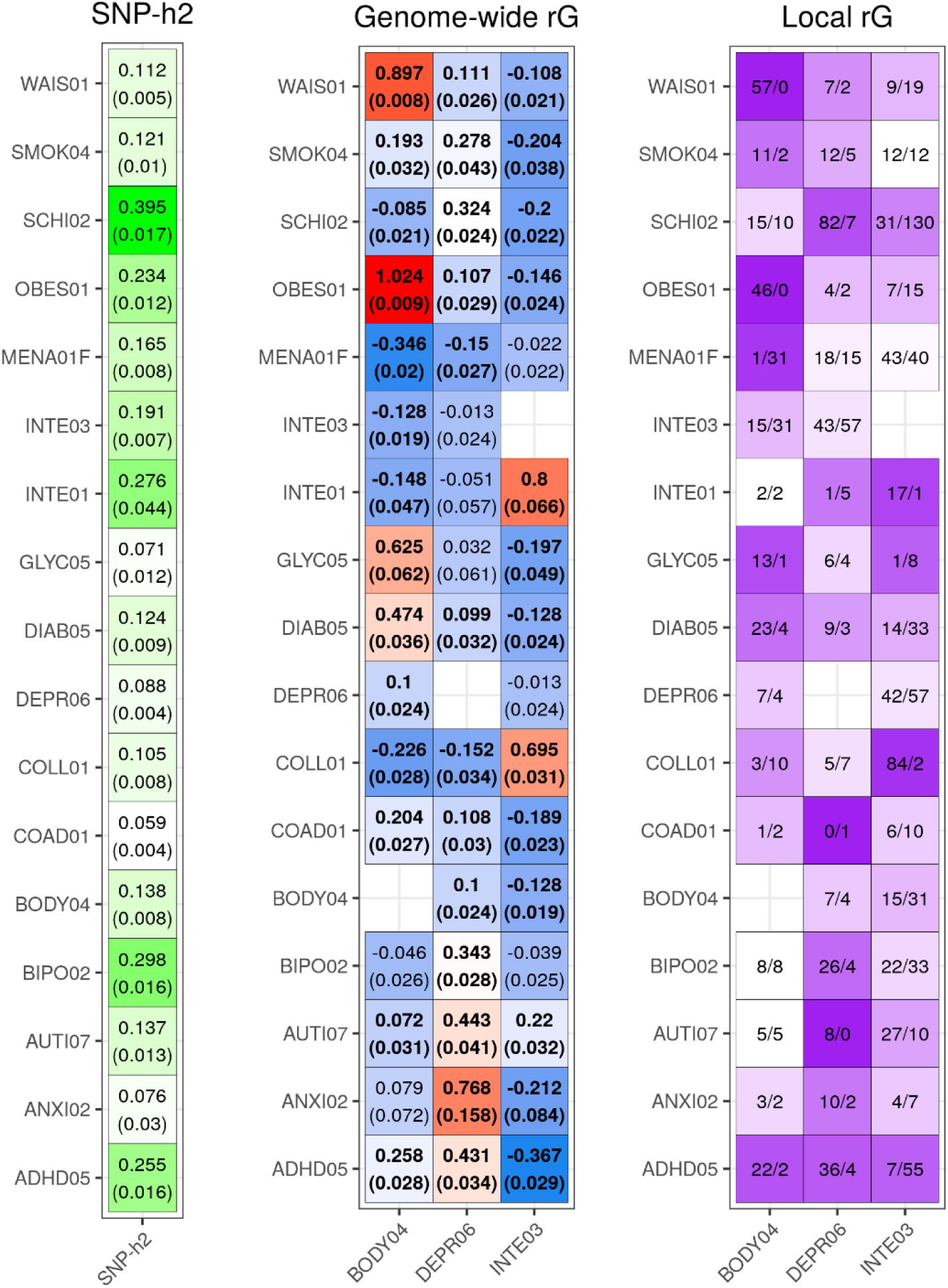
Summary of h^2^_SNP_, genome-wide r_g_, and local r_g_ for target and secondary phenotypes. h^2^_SNP_ and genome-wide r_g_ were estimated using LD score regression. h^2^_SNP_ is shown on the liability scale for binary phenotypes. The standard error of h^2^_SNP_ and genome-wide r_g_ are shown in parentheses. Significant genome-wide r_g_ estimates (p<0.05) are highlight in bold, with heat plot colours from r_g_ = 1 (red) to r_g_ = -1 (blue). Local r_g_ was estimated using LAVA. The values in the local r_g_ plot indicate the number of positive (left) and negative (right) FDR significant local genetic correlations, with the colour indicating the proportion of local genetic correlations that are in a consistent direction (deeper colour = consistent; white = discordant, with equal proportion of positive and negative correlations).

### Inclusion of PGS for secondary phenotypes improves prediction

Using an elastic net to model PGS for both the target phenotype and secondary phenotypes provided statistically significant improvements in prediction over the target phenotype PGS alone. The relative improvement in correlation between observed and predicted values for BMI, depression, and intelligence were 12.2% (*p*=1.7×10^−54^), 10.1% (*p*=7.7×10^−15^), and 0.8% (*p*=3.1×10^−12^) respectively (Figure 3). The relative improvement attained by including secondary PGS varied by the genome-wide *r*_*g*_ with the target phenotype, the *h*^2^_SNP_ and GWAS sample size of the secondary phenotype, and the predictive utility of the target PGS alone.

**Figure 3.**
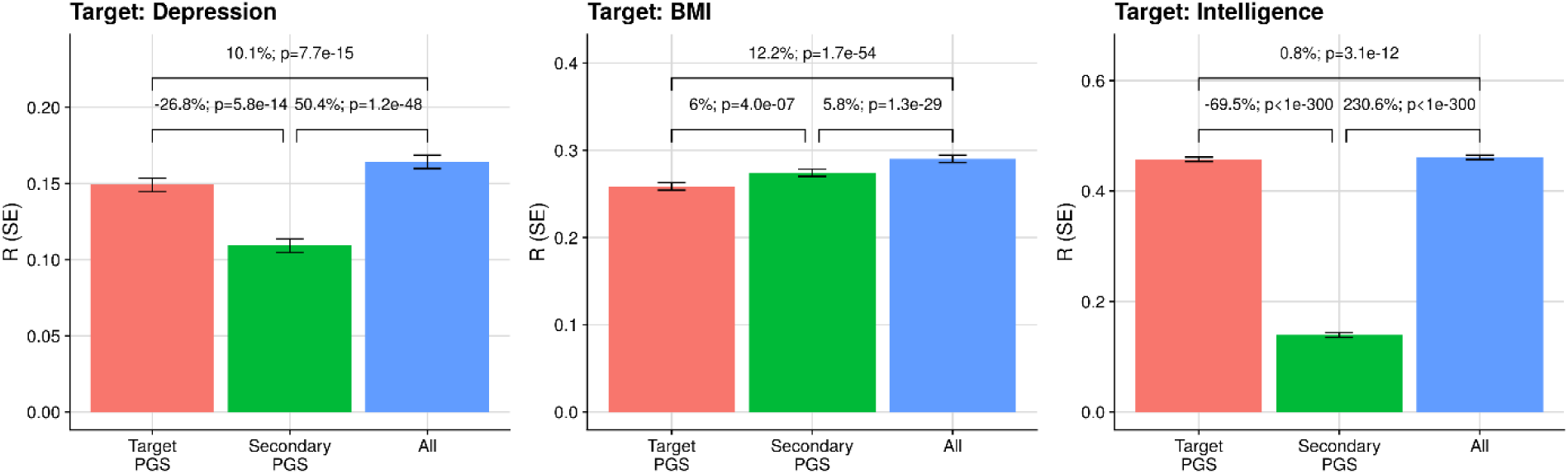
Predictive utility of models predicting depression, BMI, and intelligence. Models include target PGS only (Target PGS; red), secondary PGS only (Secondary PGS; green), and both target and secondary PGS (All); blue). Y-axis shows the Pearson correlation between predicted and observed phenotype values, with error bars indicating the standard error (note different scale by target phenotype). Pairwise comparisons of each model above the bars show the percentage difference in Pearson correlation and p-value.

### Reweighting secondary PGS by local h_SNP_^2^ and r_g_ enhances PGS across phenotypes

Here, we describe results when analysing each secondary phenotype PGS separately to determine the effect of reweighting PGS by local *h*^2^_SNP_ and *r*_*g*_. Restricting secondary phenotype PGS to variants within loci with significant local *h*^2^_SNP_ for both phenotypes or significant local *r*_*g*_ between phenotypes often led to a decrease in predictive utility compared to the genome-wide and unadjusted PGS. However, reweighting secondary phenotype PGS according to the local *r*_*g*_ with the target trait often increased the variance explained by unrestricted PGS and unweighted PGS restricted to the same loci (Figure 4).

**Figure 4.**
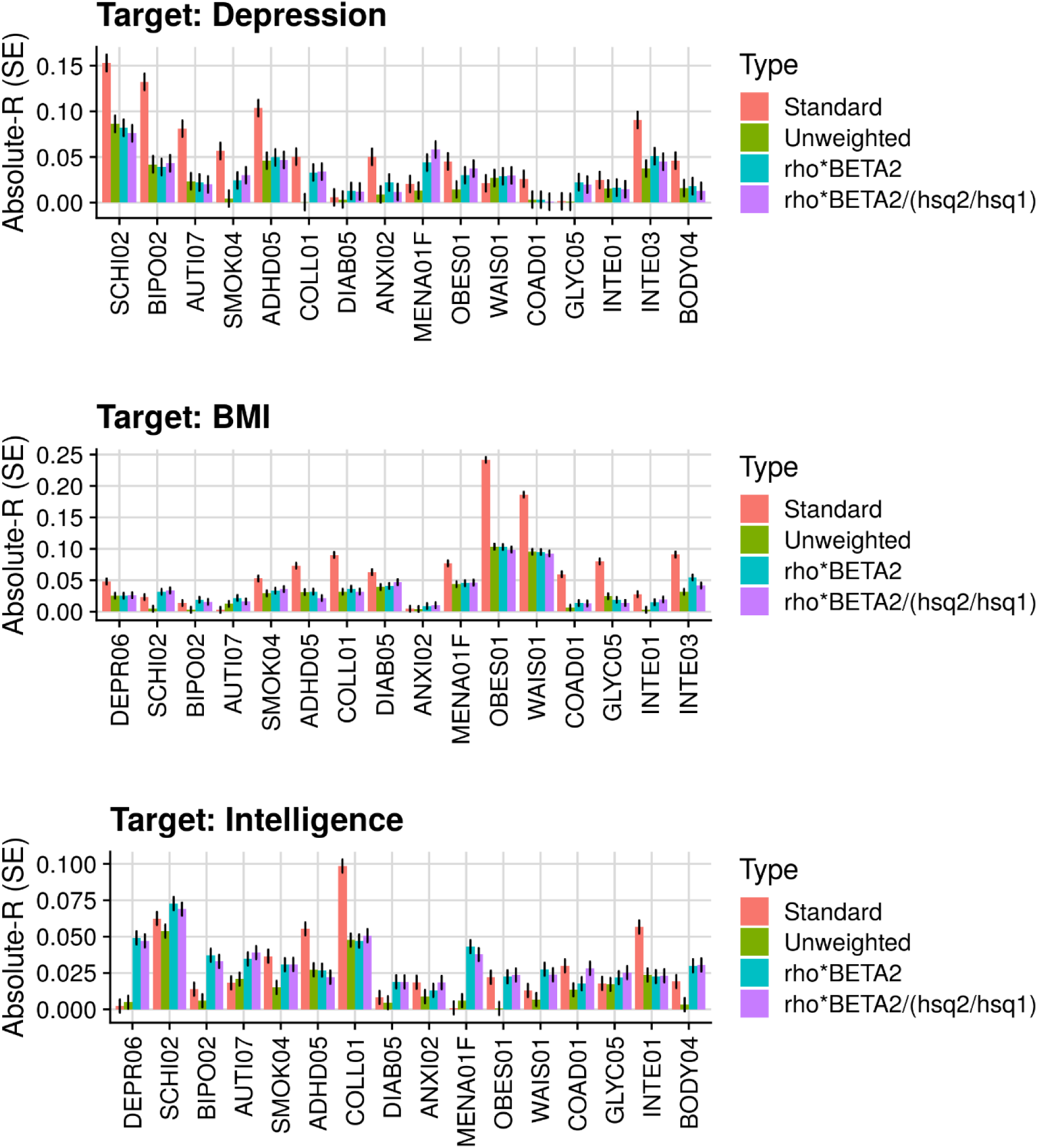
Absolute Pearson correlation between PGS and phenotypes. The ‘Standard’ PGS is an unrestricted and unadjusted PGS. All other PGS are restricted to loci with a local r_g_ p-value < 0.05 for both phenotypes. The plot compares the three SNP-weightings, unadjusted (Unweighted), adjusted for the local r_g_ (rho*BETA2), and for the local r_g_ and h^2^_SNP_ (rho*BETA2/(hsq2/hsq1)). Error bars indicate the standard error.

Modelling both reweighted secondary PGS restricted to loci with *r*_g_ *p-value* < 0.05 and unweighted secondary PGS for the remaining loci, significantly improved prediction over the standard unweighted secondary PGS alone (Figure 5). Reweighting loci by both local *r*_g_ and *h*^2^_SNP_ provided gains over standard secondary PGS of 3.8% (*p*=1.02×10^2^), 1.5% (*p*=2.5×10^−6^) and 11% (*p*=9.2×10^−14^) for depression, BMI and intelligence respectively. The relative improvement provided by the different weighting schemes varied across phenotypes, with BMI showing a significant improvement when modelling unweighted loci with *r*_g_ *p-value* < 0.05 and other loci separately, but minimal further gains when reweighting according to local *r*_g_ and *h*^2^_SNP_. In contrast, the largest improvements for depression and intelligence occurred when reweighted by local *r*_g_ and *h*^2^_SNP_.

**Figure 5.**
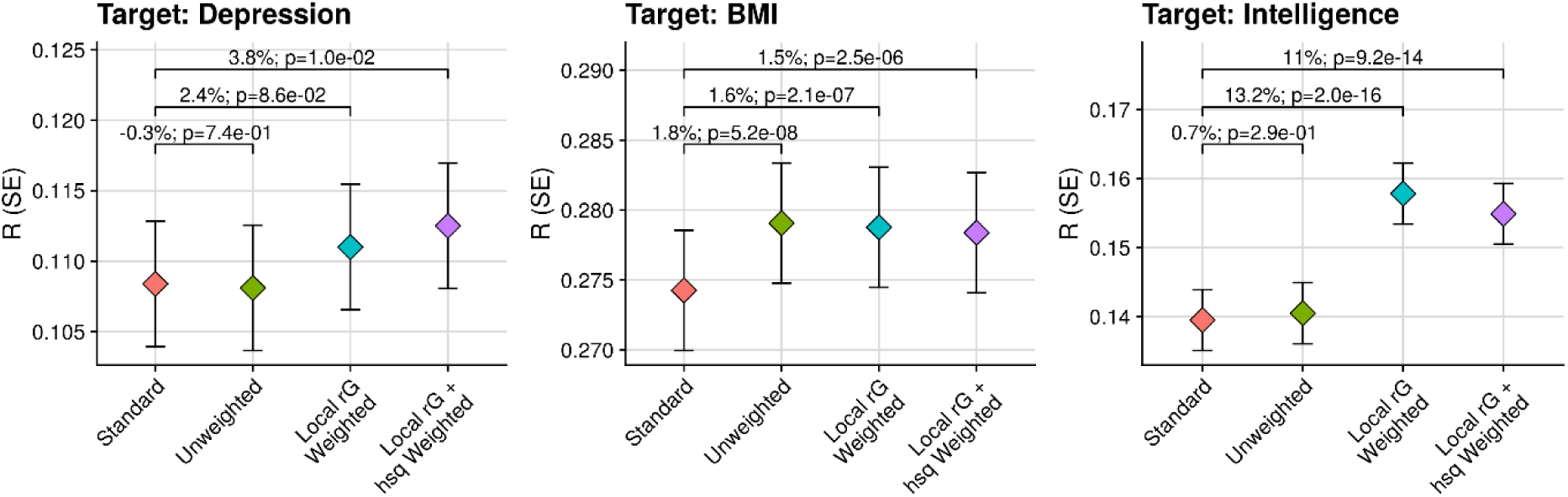
Predictive utility of models for depression, BMI, and intelligence. The ‘Standard’ model, containing standard unweighted secondary PGS, is compared to models additionally containing secondary unweighted PGS restricted to loci with non-significant local r_g_ (p-value ≥ 0.05) and PGS restricted to loci with local r_g_ p-value < 0.05 either unadjusted (Unweighted), adjusted for local rg (Local rG Weighted), or adjusted for local r_g_ and scaled by the ratio of local h^2^_SNP_ between target and secondary phenotypes (Local rG + hsq Weighted). Y-axis shows the Pearson correlation between predicted and observed phenotype values, with error bars indicating the standard error. A comparison of each model is shown above the bars, with the text first indicating the percentage difference in Pearson correlation between each model and the Baseline model, and then showing the p-value of the difference between models.

The effect of reweighting variants by local *r*_*g*_ was greater when the direction of local *r*_*g*_ estimates were less consistent across the genome (i.e., there was a mixture of positive and negative local *r*_*g*_ estimates) (Figure 6). There was little difference when restricting loci to those with nominally significant or FDR significant local genetic correlation (Figure S1).

**Figure 6.**
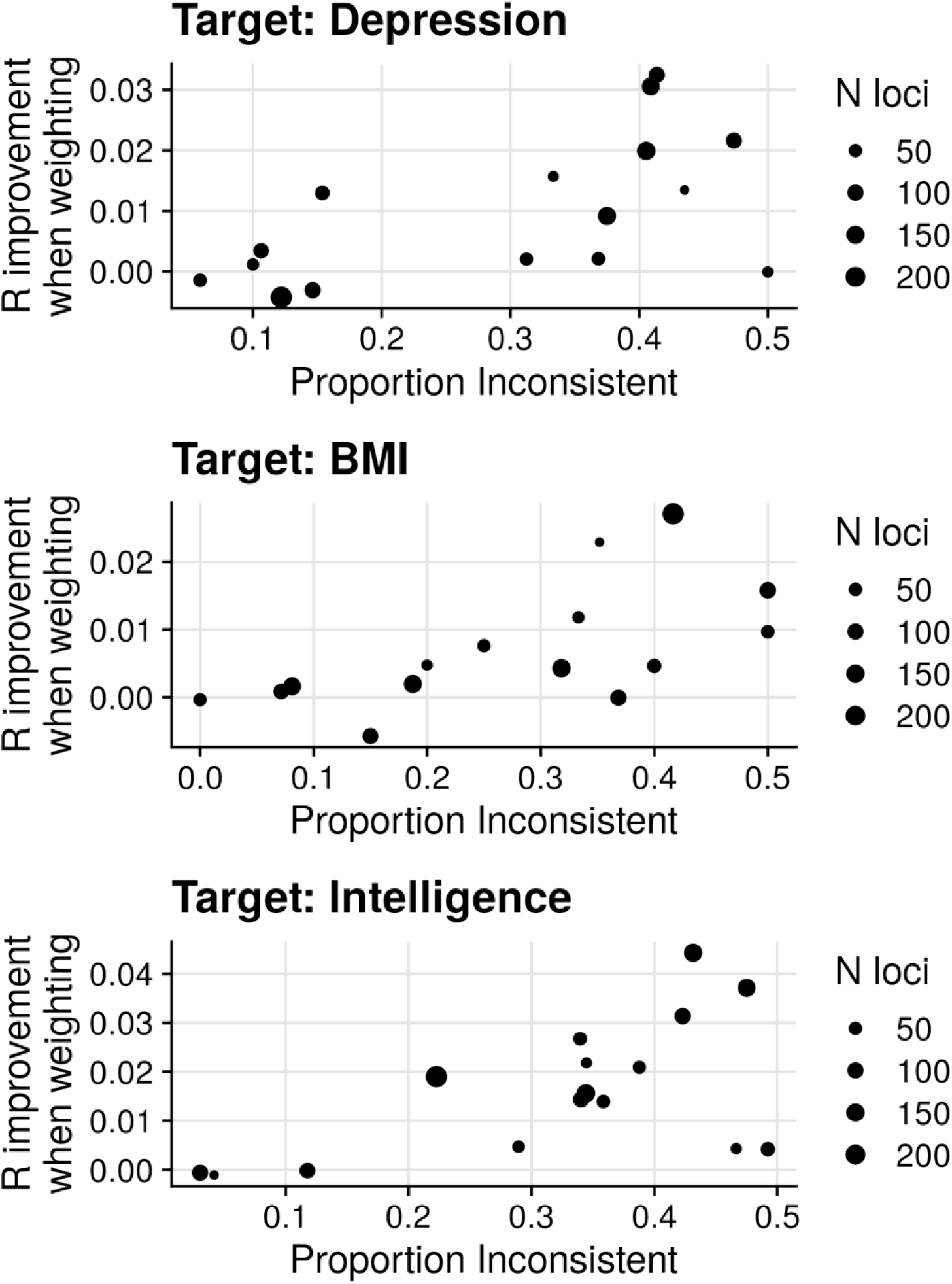
Effect of inconsistent direction in local r_g_ estimates on absolute improvement in Pearson correlation between unweighted and local r_g_ weighted PGS. All PGS are restricted to loci with nominally significant local r_g_ (p<0.05). The size of the points indicates the number of loci with nominally significant local r_g_. Proportion Inconsistent = proportion of loci with r_g_ direction opposing the most common local r_g_ direction.

### Local h^2^_SNP_ and r_g_ informed PGS can improve prediction over target and unweighted secondary PGS

We then tested whether inclusion of local *h*_SNP_^2^ and *r* informed PGS can improve prediction over models including target PGS and secondary PGS alone. The results were mixed when including secondary PGS reweighted according to local *h*^2^_SNP_ and *r*_*g*_ (Figure 7). For BMI, a relative improvement of 2.7% (*p*=2.8×10^−15^) was seen when restricting the PGS to loci with a *r*_*g*_ p-value < 0.05 without reweighting variants according to local *h*^2^_SNP_ and *r*_*g*_. Reweighting variants according to local *h*_SNP_^2^ and *r*_*g*_ provided no further gain in prediction for BMI. For Intelligence, merely restricting PGS to loci with *r*_*g*_ p-value < 0.05 provided no improvement in prediction. However, there was a nominally significant relative improvement of 0.1% (*p*=0.025) when reweighting PGS by local *r*_*g*_ estimates, and relative improvement of 0.2% (*p*=1.7×10^−3^) when reweighting PGS according to local *r*_*g*_ estimates and rescaling by differences in *h*^2^_SNP_. For depression, inclusion of the local *h*^2^_SNP_ and *r*_g_ reweighted PGS appeared to lead to overfitting, with a small decrease in prediction accuracy. There was little difference when restricting loci to those with nominally significant or FDR significant local genetic correlation (Figures S2-S4).

**Figure 7.**
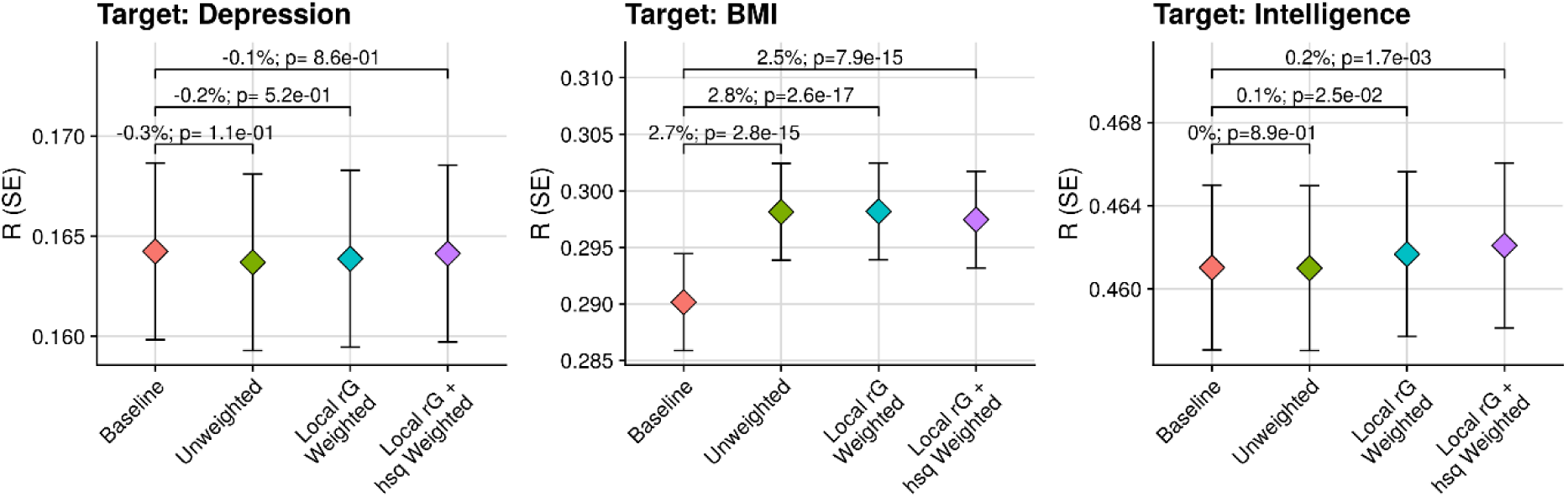
Predictive utility of models for depression, BMI, and intelligence. The ‘Baseline’ model (containing target and secondary PGS) is compared to models additionally containing secondary weighted/unweighted PGS restricted to loci with local r_g_ p-value < 0.05 either unadjusted (Unweighted), adjusted for local r_g_ (Local rG Weighted), or adjusted for local r_g_ and scaled by the ratio of local h^2^_SNP_ between target and secondary phenotypes (Local rG + hsq Weighted). Y-axis shows the Pearson correlation between predicted and observed phenotype values, with error bars indicating the standard error. A comparison of each model is shown above the bars, with the text first indicating the percentage difference in Pearson correlation between each model and the Baseline model, and then showing the p-value of the difference between models.

## Discussion

This study has evaluated a multivariate PGS approach for improving prediction of three target phenotypes in the UK Biobank sample. We initially estimate genome-wide and local heritability and genetic correlations between phenotypes using LAVA. We then evaluate the gain in target phenotype prediction by including secondary PGS derived using MegaPRS, and secondary PGS restricted and reweighted according to local heritability (*h*^2^_SNP_) and genetic correlation (*r*_*g*_ ) estimates.

Comparison of genome-wide and local estimates of *h*^2^_SNP_ and *r*_g_ demonstrates that there are often significant positive and negative local *r*_*g*_ even in the absence of significant genome-wide *r*_*g*_. This finding supports previous literature and highlights the novel insights that local methods can provide into the overlapping genetic effects between phenotypes (7, 8). Furthermore, the presence of mixed direction of local genetic correlations highlights the limitation of current multivariate PGS methods, which rely on genome-wide estimates between target and secondary phenotypes.

We then applied elastic net models containing PGS for the target phenotype and secondary phenotypes, derived using MegaPRS. This analysis showed that inclusion of PGS for secondary phenotype improved target phenotype prediction over the target phenotype PGS alone. This finding is congruent with a previous study evaluating the predictive utility of secondary PGS in the Twins Early Development Study (TEDS)(3).

To explore the utility of local *h*^2^_SNP_ and *r*_*g*_ estimates when using secondary PGS to predict the target phenotype, we first compared standard secondary PGS to secondary PGS restricted to loci with a significant *h*^2^_SNP_ for both traits and significant *r*_*g*_ between traits. In many instances the standard PGS was a better predictor of the target phenotype than *h*^2^_SNP_ and *r*_g_ restricted PGS. This highlights that locus with non-significant *h*^2^_SNP_ and *r*_*g*_ can still contribute to the variance explained by secondary PGS. This may occur due to reduced power in the target phenotype GWAS for detection of statistically significant local *h*^2^_SNP_ or *r*_*g*_.

We then tested whether leveraging local *h*^2^_SNP_ or *r*_*g*_ estimates can increase the predictive utility of secondary PGS compared to unweighted secondary PGS restricted to the same loci. As expected, based on the mixture of positive and negative local *r*_*g*_ estimates between target and secondary phenotypes, reweighting PGS according to local *r*_*g*_ estimates did often lead to an increased correlation between the secondary PGS and the target phenotype, compared to the unweighted PGS based on the same loci. Conversely, reweighting PGS by difference in local *h*^2^_SNP_ had no consistent effect on the predictive utility of the secondary PGS. These findings support the concept that accounting for local genetic correlation estimates when using GWAS/PGS for secondary phenotypes can improve target phenotype prediction. To highlight the effect of reweighting loci by *h*^2^_SNP_ or *r*_*g*_ more clearly, we evaluated the predictive utility of models containing reweighted secondary PGS restricted to loci with significant local *r*_*g*_ and unweighted secondary PGS for the remaining loci that did not have a significant local *r*_*g*_. This analysis again clearly demonstrated that reweighting secondary PGS according to local *h*_SNP_^2^ or *r*_*g*_ significantly increased the prediction of the target phenotype.

Finally, we demonstrate that inclusion of secondary PGS restricted to loci that have a significant *r*_*g*_ for both phenotypes, and reweighting PGS according to local *r*_*g*_ estimates, provides further improved target phenotype prediction over target and secondary PGS alone. However, the gain in prediction accuracy by including reweighted secondary PGS was limited, and for depression the inclusion of reweighted secondary PGS reduced prediction accuracy. This highlights a tradeoff between inclusion of additional weak predictors and the risk of overfitting, even when using large training samples and penalized regression such as elastic net. Further methodological development is required to harness the increased predictive utility of PGS reweighted according to *r*_*g*_ or *h*^2^_SNP_, whilst preserving the information in other regions of the genome.

Here we discuss several limitations of the current study and possible future directions. First, we only compare these approaches using three target phenotypes and a modest selection of GWAS summary statistics based on previously reported genome-wide *r*_*g*_ estimates. Future studies should compare methods using a wider range of target phenotypes and secondary phenotype GWAS. Second, we do not compare our approach directly with other multivariate PGS methods such as MTAG, SMTpred and GSEM. We have focused on an adaptation only for the elastic net approach for simplicity, but it is possible that adaptions of these methods to account for local *h*^2^_SNP_ and *r*_*g*_ may provide further gains in prediction. Third, we only use LAVA to estimate local genetic correlation, but other methods such as HESS are available. Fourth, local *h*^2^_SNP_ and *r*_*g*_ can only be reliably incorporated if an independent and well powered GWAS for the target phenotype is available.

In conclusion, we demonstrate that local heritability and genetic correlation can enhance the target phenotype prediction when using PGS for secondary phenotypes. We expect this more granular view of genetic overlap to be an important advance over current multivariate PGS methodology, and it should be integrated into other multivariate PGS methodologies in the future.

## URLs

- Updated GenoPred PGS methods comparison including LDAK’s MegaPRS:

https://opain.github.io/GenoPred/Determine_optimal_polygenic_scoring_approach_update

21102021.html

## Acknowledgements

The authors acknowledge use of the research computing facility at King’s College London, Rosalind (https://rosalind.kcl.ac.uk), which is delivered in partnership with the NIHR Maudsley BRC, and part-funded by capital equipment grants from the Maudsley Charity (award 980) and Guy’s & St. Thomas’ Charity (TR130505). The views expressed are those of the authors and not necessarily those of the NHS, the NIHR or the Department of Health and Social Care. We thank the research participants and employees of 23andMe, Inc for making the work regarding Depression possible.

UKB: This research was conducted under UK Biobank application 18177.

## Supplementary Figures

**Figure S1.**
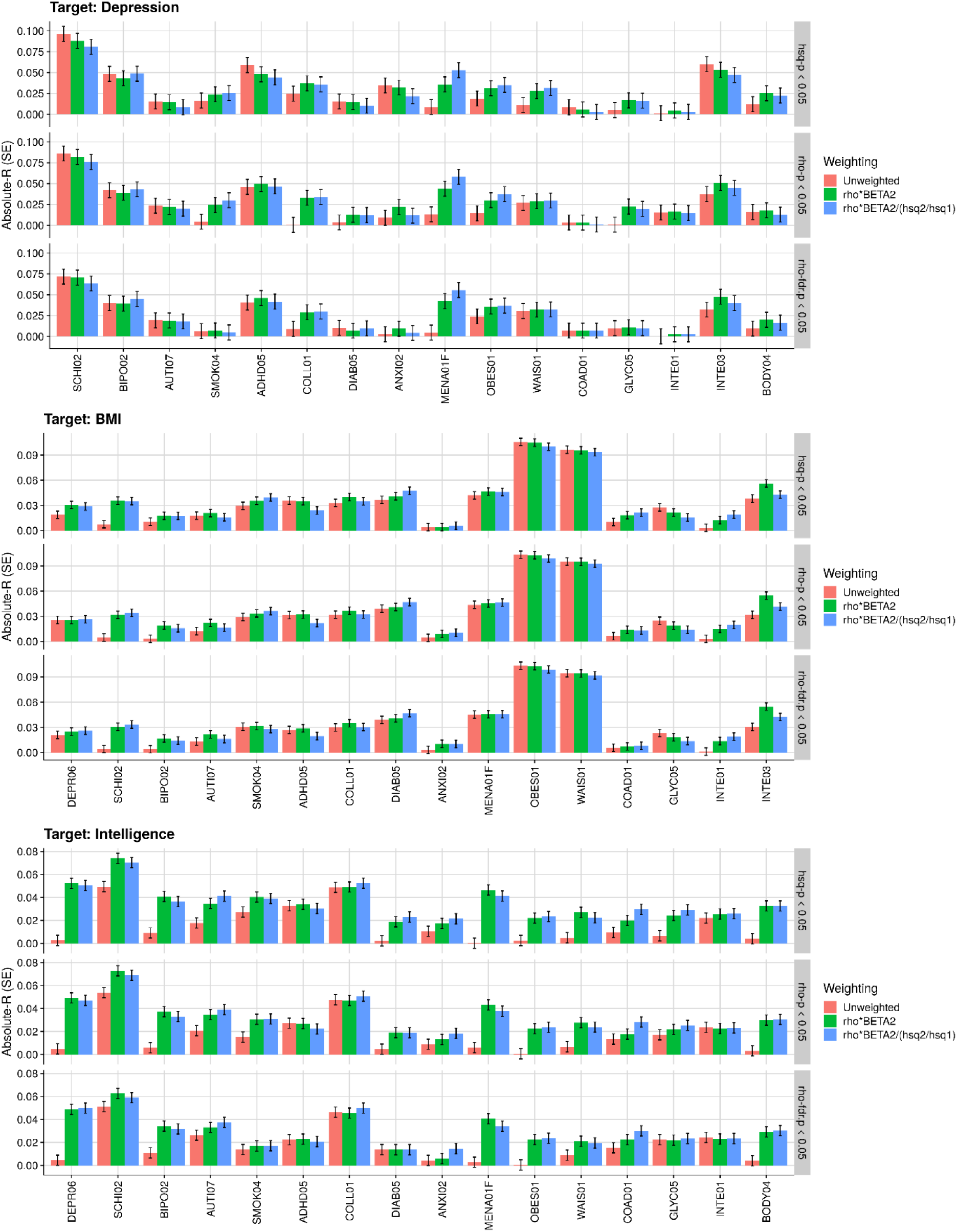
Absolute Pearson correlation between PGS and phenotypes. PGS are restricted to loci with either a local h^2^_SNP_ p-value < 0.05 for both phenotypes (hsq-p < 0.05), a nominally significant local r_g_ (rho-p < 0.05), or an FDR significant local r_g_ (rho-fdr.p < 0.05). The plot compares unadjusted PGS SNP-weights (Unweighted), PGS SNP-weights adjusted for the local r_g_ (rho*BETA2), and PGS SNP-weights adjusted for the local r_g_ and h^2^_SNP_ (rho*BETA2/(hsq2/hsq1)). Pearson correlation standard errors are shown in parentheses.

**Figure S2.**
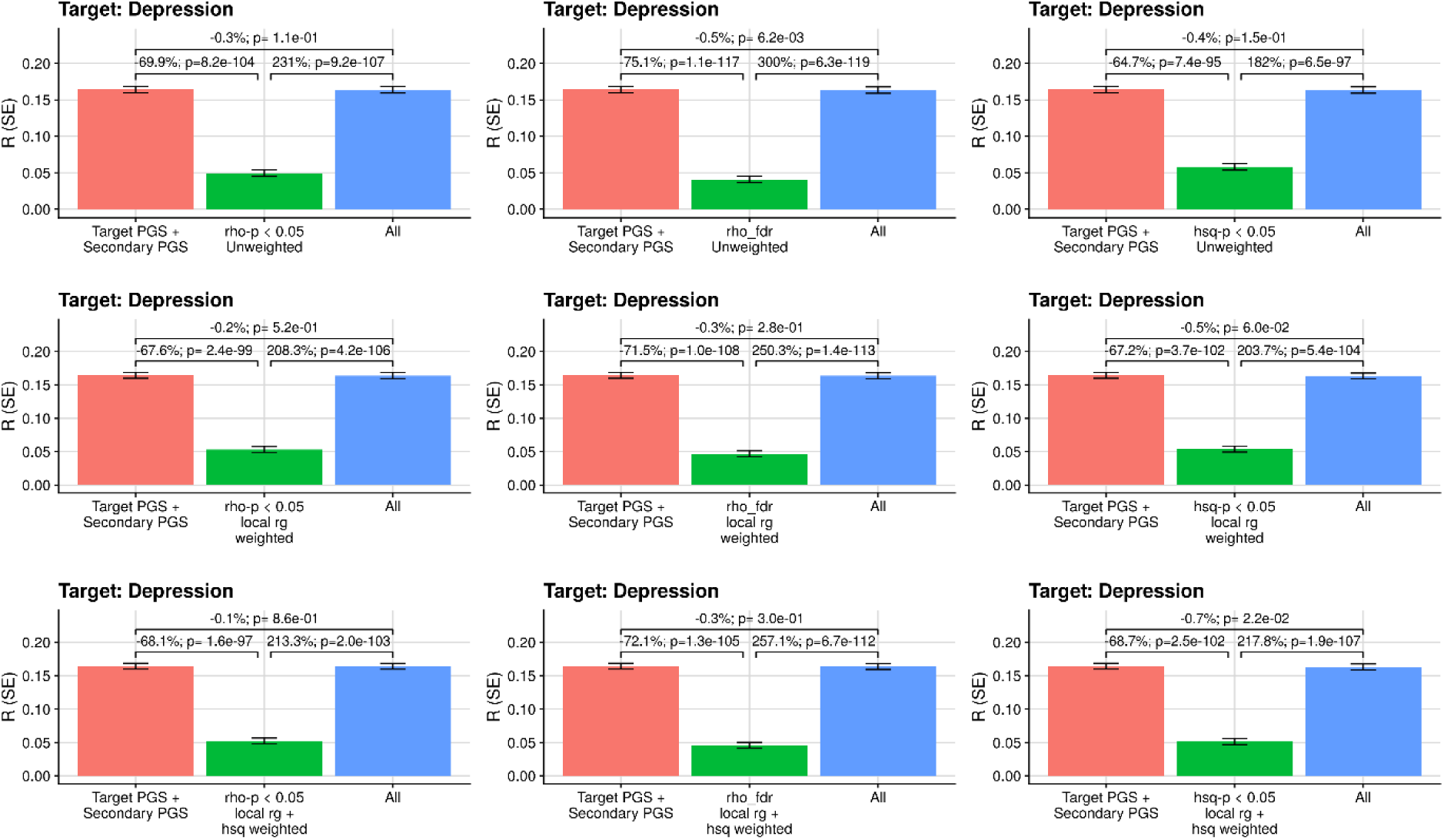
Predictive utility of models predicting depression. Models include either target and secondary PGS, secondary weighted/unweighted and restricted PGS, or both target and secondary PGS, and secondary weighted/unweighted and restricted PGS (All). The secondary PGS are either restricted to loci with a local h^2^_SNP_ p-value < 0.05 (hsq-p < 0.05), a nominally significant local r_g_ (rho-p < 0.05), or an FDR significant local r_g_ (rho-fdr.p < 0.05), and were either unadjusted (Unweighted), adjusted for local r_g_ (local rg weighted), or adjusted for local r_g_ and scaled by the difference in local h^2^_SNP_ (local rg + hsq weighted). Y-axis shows the Pearson correlation between predicted and observed phenotype values, with error bars indicating the standard error. A comparison of each model is shown above the bars, with the text first indicating the percentage difference in Pearson correlation between the left and right bar, and then showing the p-value of the different between models.

**Figure S3.**
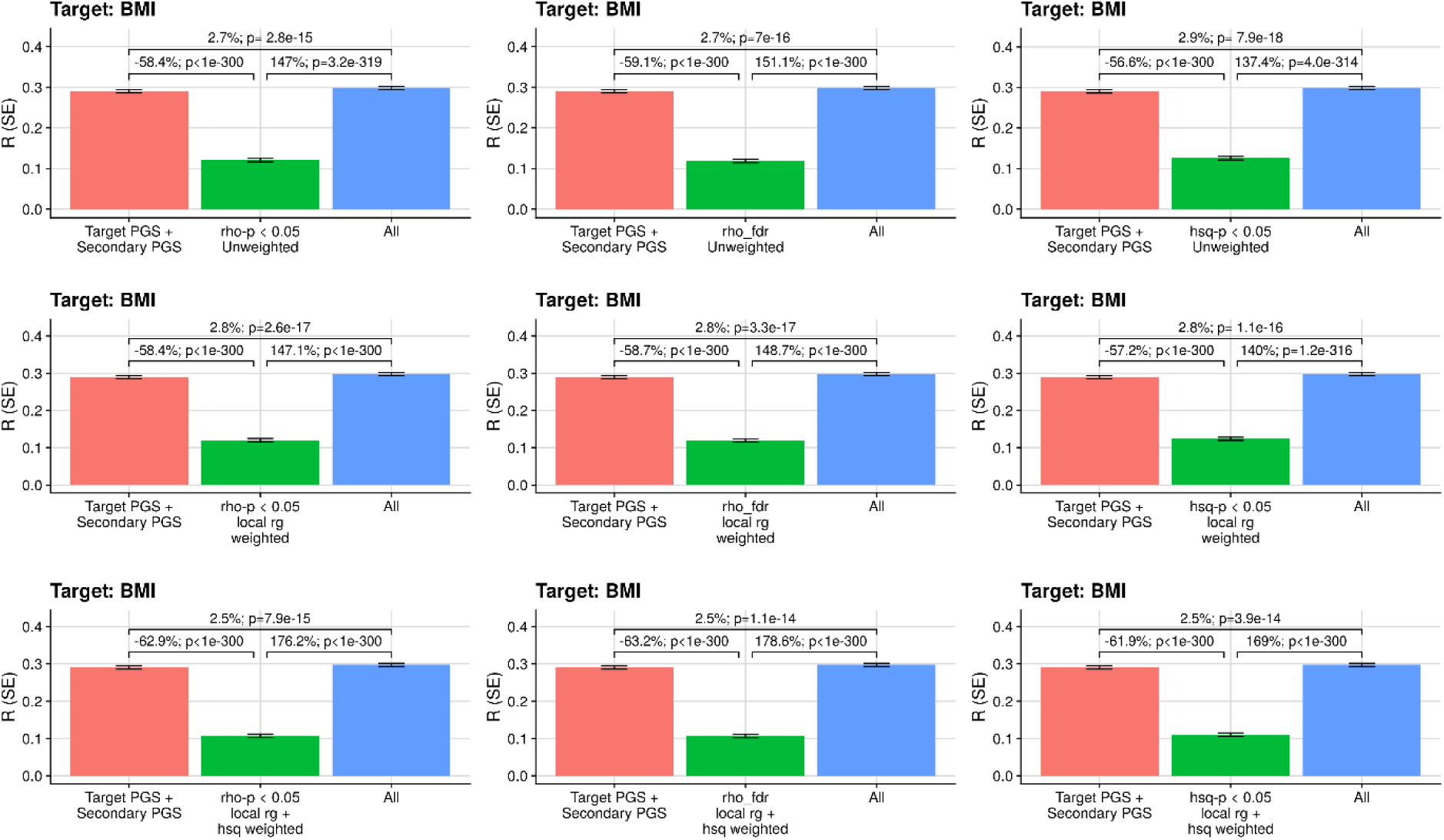
Predictive utility of models predicting BMI. Models include either target and secondary PGS, secondary weighted/unweighted and restricted PGS, or both target and secondary PGS, and secondary weighted/unweighted and restricted PGS (All). The secondary PGS are either restricted to loci with a local h^2^_SNP_ p-value < 0.05 (hsq-p < 0.05), a nominally significant local r_g_ (rho-p < 0.05), or an FDR significant local r_g_ (rho-fdr.p < 0.05), and were either unadjusted (Unweighted), adjusted for local r_g_ (local rg weighted), or adjusted for local r_g_ and scaled by the difference in local h^2^_SNP_ (local rg + hsq weighted). Y-axis shows the Pearson correlation between predicted and observed phenotype values, with error bars indicating the standard error. A comparison of each model is shown above the bars, with the text first indicating the percentage difference in Pearson correlation between the left and right bar, and then showing the p-value of the different between models.

**Figure S4.**
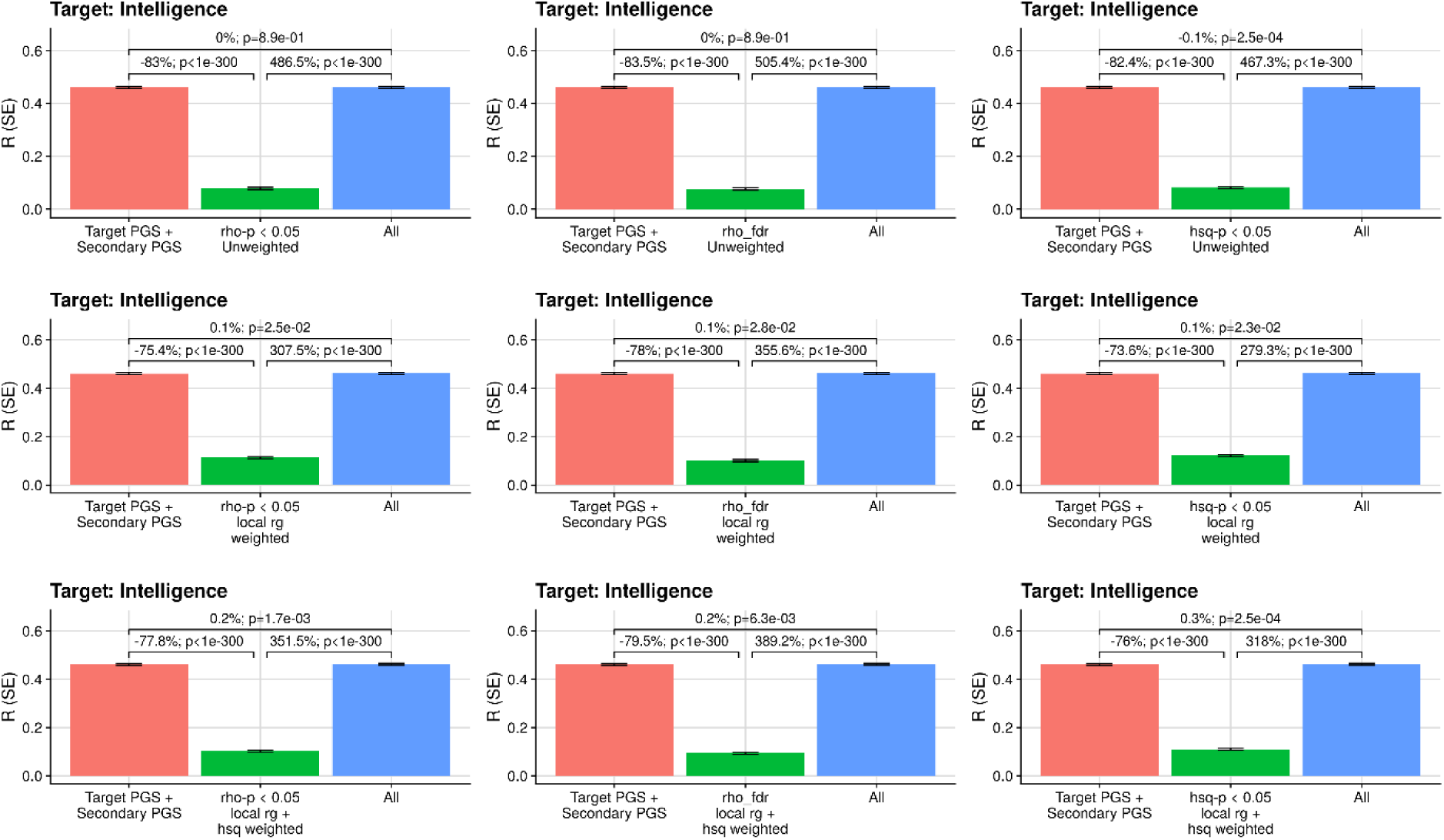
Predictive utility of models predicting intelligence. Models include either target and secondary PGS, secondary weighted/unweighted and restricted PGS, or both target and secondary PGS, and secondary weighted/unweighted and restricted PGS (All). The secondary PGS are either restricted to loci with a local h^2^_SNP_ p-value < 0.05 (hsq-p < 0.05), a nominally significant local r_g_ (rho-p < 0.05), or an FDR significant local r_g_ (rho-fdr.p < 0.05), and were either unadjusted (Unweighted), adjusted for local r_g_ (local rg weighted), or adjusted for local r_g_ and scaled by the difference in local h^2^_SNP_ (local rg + hsq weighted). Y-axis shows the Pearson correlation between predicted and observed phenotype values, with error bars indicating the standard error. A comparison of each model is shown above the bars, with the text first indicating the percentage difference in Pearson correlation between the left and right bar, and then showing the p-value of the different between models.

